# Allosteric regulation of the proteasome’s catalytic sites by the proteasome activator PA28γ /REGγ

**DOI:** 10.1101/2021.12.09.471496

**Authors:** Taylor A Thomas, David M. Smith

## Abstract

Proteasome Activator 28γ (PA28γ) is a member of the 11S family of proteasomal regulators that is constitutively expressed in the nucleus and is implicated in certain cancers, lupus, rheumatoid arthritis, and Poly-glutamine neurodegenerative diseases. However, how PA28γ functions in protein degradation remains unclear. Though PA28γ’s mechanism has been investigated for some time, many alternative hypotheses have not been tested: e.g. 1) substrate selection, 2) allosteric upregulation of the Trypsin-like catalytic site, 3) allosteric inhibition of the Chymotrypsin- and Caspase-like catalytic sites, 4) conversion of the Chymotrypsin- or Caspase-like sites to new Trypsin-like catalytic sites, and 5) gate-opening in combination with these. The purpose of this study was to conclusively determine how PA28γ regulates proteasome function. Here, we rigorously and definitively show that PA28γ uses an allosteric mechanism to upregulate the proteolytic activity of the 20S proteasome’s Trypsin-like catalytic site. Using a constitutively open channel proteasome, we were able to dissociate gating affects from catalytic affects demonstrating that the PA28γ-increases the affinity (K_m_) and V_max_ for Trypsin-like peptide substrates. Mutagenesis of PA28γ also reveals that it does not select for (i.e. filter) peptide substrates, and does not change the specificity of the other active sites to trypsin-like. Further, using Cryo-EM we were able to visualize the C7 symmetric PA28γ-20S proteasome complex at 4.4Å validating it’s expected 11S-like quaternary structure and proteasome binding mode. The results of this study provide unambiguous evidence that PA28γ functions by allosterically upregulating the T-L like site in the 20S proteasome.

**Significance Statement:** This study rigorously demonstrates that PA28g allosterically activates the b-2 proteolytic site of the 20S proteasome directly without affecting 20S gating. Further, we generated the first human 11S-human 20S proteasome cryo-EM structure of the PA28g-20S complex showing that, despite it’s different affects on 20S activity, it has a similar quaternary structure as the other 11S family members. The significance of these findings is paramount as the b-2 site is responsible for post-basic cleavage and suggests that PA28g is specialized to degrade positively charged DNA binding proteins. Further, b-2 upregulation via PA28g could provide a protective effect against poly-glutamine expanded proteins, like Huntingtin. This work provides a framework for PA28g drug development to treat PA28g addicted cancers and Huntington’s Disease.

## Introduction

Proteasome Activator 28γ (PA28γ, also known as Ki antigen, REGγ) is a proteasomal activator in the 11S family that is implicated in several cancers^1–3^ and rheumatoid arthritis^4^ where it is found to be overexpressed. Under normal physiological conditions, PA28γ is constitutively expressed across all tissues^5^ and is localized to the nucleus, but not within the nucleolus^6^. Interestingly, expression of PA28γ has been shown to enhance survival in an *in vitro* model of Huntingdon’s Disease^7^, and gene therapy of PA28γ improves motor coordination in a murine Huntington’s Disease model, YAC128^8^. However, in a cell model of spinal and bulbar muscular atrophy (SMBA), which is caused by polyglutamine (PolyQ) expansion in the androgen receptor (AR), PA28γ had an adverse effect on AR aggregation in a proteasome-independent manner^9^. Despite these physiological and cell biological findings regarding PA28γ’s biological roles, the molecular mechanism of how PA28γ stimulates protein degradation by the proteasome remains a mystery.

The 11S family of proteasomal activators extends across all multicellular eukarya, and are often interchangeably referred to as PA26/PA28 or REG. This class of activators are all heptameric complexes, ATP-independent and do not contain unfolding or forced translocation activity^10^. The 11S family is an expanding class of proteasomal activators found in a variety of species such as *Trypanosoma brucei* (PA26)^11^, *Drosophila*^12^, *Plasmodium falciparum* (*Pf*)^13^, ticks^14^, and mammals^10^. In mammals, there are 3 homologs of PA28 within the 11S family: alpha (α), beta (β), and gamma (γ); PA28α and PA28β form an asymmetric heteroheptameric complex known as PA28αβ^15^. PA28αβ expression is regulated by interferon-γ and has been identified to play a functional role in Major Histocompatibility (MHC) Class I antigen presentation^16,17^. In contrast, PA28γ is a homoheptameric complex, that does not form a complex with PA28α or PA28β^10^. As described above, PA28γ is implicated in a variety of disease states but its physiologic role in nuclear proteostasis and how it regulates protein degradation through the proteasome remains unclear. Further, PA26 and PA28αβ complexes have been structurally characterized using X-ray crystallography, and the PA28αβ-immunoproteasome complex has been solved using cryo-EM, but the structure of PA28γ complex remains unknown.

The eukaryotic 20S proteasome is a compartmentalized protease comprised of four, heteroheptameric rings in an α, β, β, α arrangement^18,19^. The β rings each contain 3 active sites and sequesters their protease activity to the hollow interior of the 20S^20^. Each active site has unique specificity for amino acid side chains (i.e., β5 – chymotrypsin-like (CT-L), β2 - trypsin-like (T-L), and β1 - caspase-like (C-L))^21^. An α-ring flanks the β rings on either side, and the N-termini of the α-subunits fold over the substrate entry pore to form a barrier to the degradation chamber. This barrier is called the proteasomal gate and it protects the cell from unregulated protein degradation^22^. Regulating the proteasomal gate and the precise degradation of cellular substrates is managed by a variety of proteins or protein complexes called proteasomal activators.

Proteasomal activators are highly specialized and have unique functions. Proteasomal ATPases (i.e., regulatory particle or 19S) utilize ATP to recognize, unfold, and translocate substrates into the degradation chamber^23,24^. ATP-independent proteasome activators include PA200/Blm10, PI31, and 11S, each with their own unique regulatory mechanisms. Our current understanding of the 11S-proteasome complex mechanism is based on a novel crystallography study of PA26 in complex with the yeast and archaeal proteasomes, which not only contributed to our understanding of the activation mechanism but demonstrated the 11S family’s capability to activate the proteasome across multiple species^25^. Whitby et. al. revealed that PA26 regulates substrate entry via opening the gate of the 20S proteasome. Opening the 20S gate allows peptide substrates to freely defuse through the center of the 11S regulators pore and into the 20S catalytic chamber, where they are degraded. Interestingly a mechanism for stimulating unfolded-protein degradation has not been worked out but is expected to be similar. Recently, two cryo-EM studies substantiated similar mechanisms of proteasomal regulation for PA28αβ and the immunoproteasome, and the *Pf*PA28-20S complex, indicating that PA26, PA28αβ, and *Pf*PA28 use similar mechanisms to regulate proteasome function^13,26^. In contrast, since its characterization in the 1990s, PA28γ’s effect on the proteasome has been under contention. Initial studies of PA28γ reported conflicting observations regarding how PA28γ alters 20S proteasome activity. It was determined that different PA28γ purification strategies were the reason for these different observations. Wilk et. al used ammonium precipitation to purify PA28γ and concluded that it had 20S gate-opening activity and did not observe activation of a specific proteolytic activities^27^. However, alternative purification strategies and extracting PA28γ from a variety of cell lines lead to the conclusion that endogenous PA28γ specifically upregulates the cleavage of T-L peptides ^28^. While it was suggested that this upregulation was due to catalytic activation of the T-L like sites in the 20S, the mechanism was never determined, and several potential alternative mechanisms were never ruled out. Moreover, a recent cryo-EM structure of human PA200 and human 20S proteasome revealed a unique mechanism of T-L proteolytic activation through switching of the β5 S1 pocket^29^, which effectively turns the CT-L site into a T-L site, thus providing another alternative mechanism by which T-L peptide hydrolysis could be upregulated by PA28γ. The discovery of this novel mechanism indicates the need to understand the precise mechanism of proteasomal activation by PA28γ. Five alternative hypothesis could explain how PA28γ selectively stimulates T-L peptide hydrolysis: 1) specific substrate selection, 2) allosteric upregulation of β2 catalytic site (T-L site), 3) allosteric inhibition of only the β1 and β5 catalytic sites, 4) alteration of β1 or β5 substrate specificity to more T-L like, or 5) proteasomal gate opening in combination with one of the previous hypothesis^30,31^. However, in previous years, experimentally decoupling the effects PA28γ has on either gate opening, or the catalytic sites would have been very difficult to test as using the constitutive 20S proteasome complicates the analysis due to the potential of gating effects.

The present study was undertaken to rigorously investigate which of these mechanisms PA28γ employs to upregulate T-L peptide substrate proteolysis by the proteasome. Understanding the mechanism of this proteolytic stimulation would be important for understanding changes to proteostasis during diseases, such as in cancers where PA28γ expression is upregulated or during Huntingdon’s Disease, where the proteasome is inefficient at degrading proteins with poly-glutamine (Poly-Q) expanded repeats, which are primarily degraded by the T-L site^32^. To investigate the mechanism of PA28γ’s proteasomal regulation, we employed the α3ΔN-20S or “open-channel” proteasome. α3ΔN is a mutant proteasome species that has an N-terminal, 10-residue truncation of the α3 subunit. The α3 N-termini stabilizes the closed state of the proteasome by interacting with α2 and α4 in the 20S pore. Thus, the α3N-termini deletion produces a proteasome that has a constitutively open channel^22^. Combining the α3ΔN-20S with PA28γ allows us to mechanistically decouple proteasomal gate-opening from potential allosteric activation of a protease site. This combined with other protein engineering tools allows for the rigorous determination of mechanisms that PA28γ employs to regulate substrate degradation via the proteasome.

## Results

### WT PA28γ demonstrates upregulation of the β2 active site

Before specifically asking which hypothesis PA28γ uses to activate 20S function, we first sought to validate the activity of our purified recombinant PA28γ, as previous groups indicated different PA28γ functions that were deemed to be dependent on the specific purification strategy employed^27,28^. As controls, we also purified recombinant PA28αβ, which is known to induce gate-opening, and a mutant of PA28γ that has been reported to switch its activity. This prior study showed that a lysine mutation on the 3^rd^ helix of PA28γ, K188 to E (PA28γ-K188E), switched PA28γ’s activation activity from T-L activating to proteasomal gate-opening, thus causing it to function like PA28αβ^30^ (Fig 1A). Our recombinant human PA28γ in the presence of human 20S proteasome (H20S) induces an almost 20-fold increase in proteolytic activity for the T-L proteolytic site specific peptide substrate (RLR-AMC) when compared to H20S only controls (Fig 1C), as was similarly reported in Gao et al^28^. However, under the same conditions PA28αβ, and PA28γ-K188E stimulated substantial proteolytic activity for all peptide substrates, rather than only T-L activity (Fig 1C). These results replicate past observations that the PA28γ-K188E mutation alters PA28γ activity and likely switches it to a gate-opening form (similar to PA28αβ). This is because gate opening is expected to stimulate the degradation of all peptides regardless of their proteolytic site specificity, as it should freely allow the entry of all peptide substrates. These results clearly demonstrate that PA28γ stimulates the proteasome differently than PA28αβ, and that the K188E mutation alters how it stimulates 20S function. These results can be explained by several possible mechanisms used by PA28γ that are listed in the introduction. It is also not clear from these results alone if WT PA28γ can function via both gate-opening and T-L like activation or if these mechanisms could function mutually exclusively in WT PA28γ.

**Figure 1.**
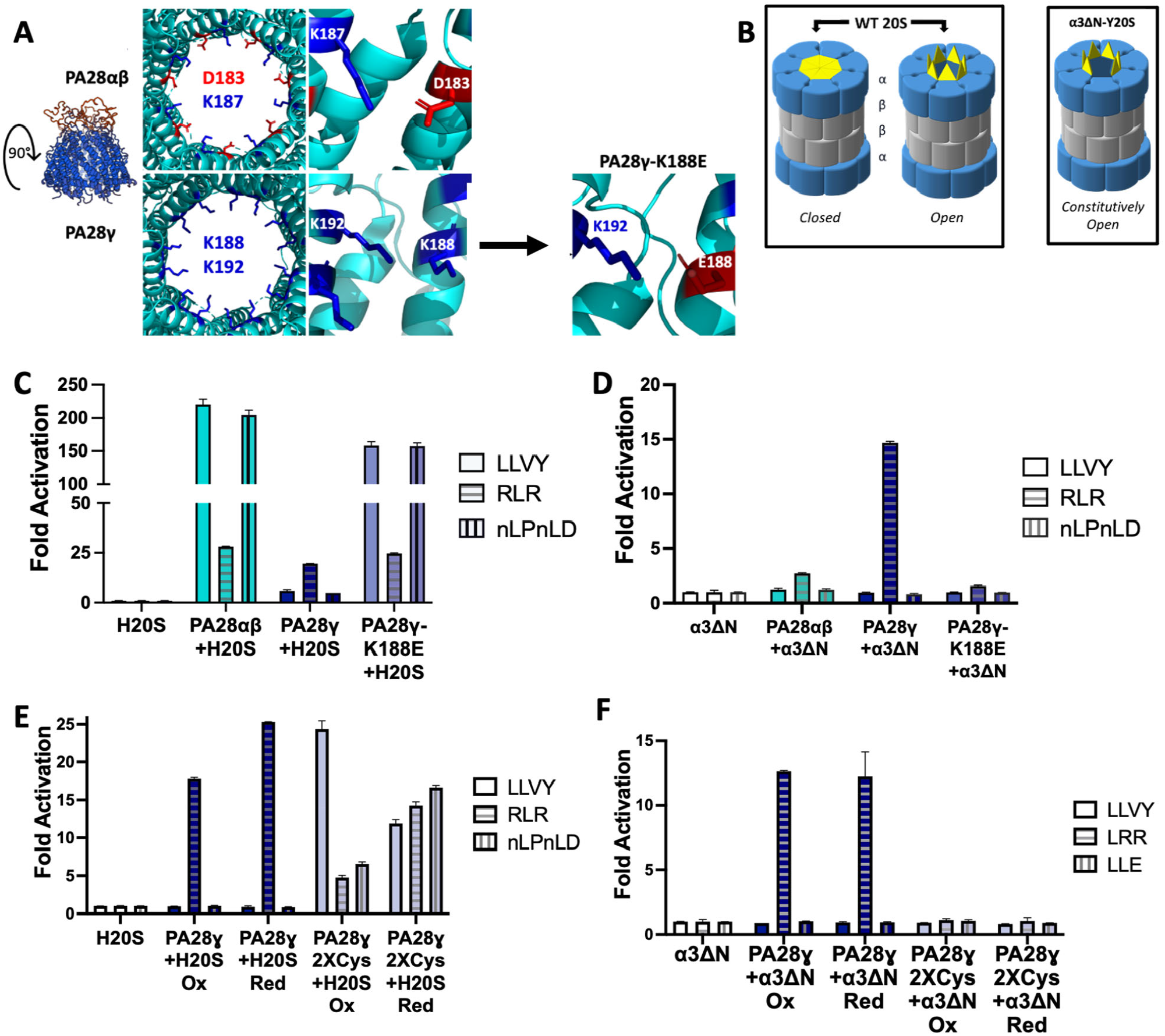
PA28γ stimulation of T-L activity does not require 20S gate-opening. A) Substrate entry pore of PA28αβ (PDB: 5MX5) alternates positive and negative charges around the ring, whereas PA28γ contains a positive substrate entry pore. PA28γ mutant, PA28γ-K188E, was generated to create a pore more like PA28αβ (PA28γ structures based on modified PDB: 5MX5). B) The WT 20S proteasome can fluctuate between a closed and open state due to thermodynamic flux, the closed state protects substrates from non-specific entry. The constitutively active, open channel α3ΔN-Y20S-proteasome has an N-terminal truncation mutation that causes the gate to remain open and allows non-specific substrate entry. C) Purified H20S proteasome (1 nM) activity was measured for all three proteolytic sites (RFU/min) in the presence of recombinant PA28αβ (50nM), PA28γ or PA28γ-K188E (62.5nM). D) Purified constitutively open-gate α3ΔN-Y20S (0.1nM) activity was measured for all three proteolytic sites (RFU/min) in the presence of recombinant PA28αβ (50nM), PA28γ or PA28γ-K188E (62.5nM). E) H20S (1nM) activity was measured for all three proteolytic sites (RFU/min) in the presence of PA28γ or PA28γ2XCys (62.5nM) under reducing and oxidizing conditions. F) Purified constitutively open-gate α3ΔN-Y20S (0.1nM) activity was measured for all three proteolytic sites (RFU/min) in the presence of recombinant PA28γ or PA28γ2XCys (62.5nM) under reducing and oxidizing conditions. Experiments were controlled for using buffer identical to the proteasomal activator. Results are the mean of at least three independent experiments performed in triplicate (error bars represent SD) normalized to the H20S or α3ΔN-Y20S control.

To test whether PA28γ directly upregulates the T-L proteolytic site or performs substrate-selective gate opening, we purified α3ΔN proteasomes from yeast (α3ΔN-Y20S). α3ΔN-Y20S has an N-terminal truncation mutation to the α3 subunit of the Y20S that alleviates the closed gate conformation induced by the N-termini of the WT Y20S α-subunits^33^ (Fig 1B). Importantly, WT yeast 20S respond similarly to activation by PA28αβ, and PA28γ as do human 20S (Supplemental Fig 1). The α3ΔN-Y20S thus mechanistically decouples gating effects from proteolytic site effects allowing us to unambiguously determine if PA28γ alters proteolytic site activity in the absence of gating contributions. We hypothesized that if an 11S family member’s proteasomal regulatory mechanism was to open the gate, there would be no change from the α3ΔN-Y20S control, as the proteasome is already constitutively active (ie. peptide substrates are free to defuse into the 20S without hinderance). However, if the activator’s regulatory mechanism included allosteric changes to the proteolytic sites, this could be clearly observed enzymatically in the absence of a functioning gate. Our results demonstrate that there were no changes to the fold activation of α3ΔN-Y20S in the presence of PA28αβ and PA28γ-K188E (Fig 1D), which demonstrates and confirms that PA28αβ and PA28γ-K188E indeed function to open the proteasomal gate allowing peptide to diffuse in, and do not themselves affect the activity of proteolytic sites. However, PA28γ induced ∼15-fold activation of the α3ΔN-Y20S in the presence of the T-L substrate RLR-AMC, when compared to the α3ΔN-Y20S controls, and did not inhibit or alter the degradation of CT-L or C-L substrates (LLVY-AMC and nLPnLD-AMC) (Fig 1D). It is important to note that the cleavage rates of the peptide substrates vary between each peptide, and that the T-L peptide, RLR-AMC has the lowest V_max_ when compared to the CT-L and C-L peptides^34^. These differences will be further analyzed below and in the discussion. These results demonstrate that PA28γ activates T-L activity of the proteasome without requiring the induction of gate-opening, but further alternative mechanisms must be ruled out to determine if this activation is catalytically allosteric. These results also conclusively demonstrate that a single point mutation is indeed capable to switch PA28γ from a T-L activation state to gate-opening state, and that these two states are mutually exclusive, as K188E mutant could not activate the α3ΔN-Y20S.

Disulfide crosslinking has been established to be a useful tool in studying proteasomal activators’ structure and function, particularly for homomers, such as the archaeal proteasomal AAA+ ATPase, PAN^35^. Briefly, cysteine point mutations are strategically induced into the protein subunit that will interact with the point mutation in the neighboring subunits. We hypothesized that if the two interacting lysine residues on helix 3 are important for maintaining PA28γ T-L activity, and if the gate-opening state is the result of the lysine-glutamate salt bridge interactions in PA28γ-K188E and PA28αβ then inducing two hydrophobic residues, like cysteine, would also change the activity of PA28γ from T-L stimulating to gate opening. We adopted the previous approach for PA28γ and induced two cysteine point mutations at the K188 and K192 positions within helix 3, which will further be called PA28γ2XCys, similar to the K188 to E mutation (Fig 1A). When tested in a proteasome activity assay using WT H20S or α3ΔN-Y20S, the results reveal that PA28γ2XCys has similar activity patterns to that of PA28γ-K188E or PA28αβ and opens the proteasomal gate without inducing T-L activation (Fig 5B and C). As cysteine residues can form disulfide bonds, we tested these mutants under both reducing, non-crosslinked, and oxidizing, crosslinked, conditions. Under both conditions, PA28γ2XCys stimulated all three activities in the WT 20S, but did not stimulate the open channel 20S indicating that it induces only gate opening. One caveat is that PA28γ2XCys did not stimulate gate-opening as well as did PA28γK1888E, presumably due to a less efficient conformational state induced by changing hydrophilic pore residues to hydrophobic ones. Nevertheless, these results further demonstrate that pore stabilizing mutations in PA28γ, whether salt-bridge, hydrophobic, or covalent in nature can switch PA28γ from an allosteric activating state to a gate-opening state.

### PA28γ does not alter the specificity of the CT-L or C-L proteolytic sites to upregulate T-L activity

While intuitively one would think that an increase in T-L like activity is likely due to an increase in the proteolytic activity of the T-L proteolytic site, this may not necessarily be the case. A recent cryo-EM study of the human PA200-20S complex determined that PA200’s upregulation of T-L activity is the result of conformational changes to the S1 pocket of β5 which causes β5 to change from CT-L activity to T-L activity like β2^29^. Therefore, we asked if it was possible PA28γ was using a similar mechanism to upregulate T-L activity. We hypothesized that if PA28γ caused allosteric changes to the β5 S1 binding pocket to increase T-L activity, then inhibiting the β5 CT-L site prior to binding of PA28γ, would block PA28γ ability to stimulate T-L activity. Alternatively, if PA28γ allosterically upregulates β2 or is selective for T-L peptide substrates, we should see loss of that activity in the presence of a β2 inhibitor. Therefor we employed the use of proteasome inhibitors epoxomicin (irreversible inhibitor) and leupeptin that specifically target β5 and β2, respectively^36^. We began the experiment by incubating the H20S with the inhibitor prior to running the proteasome activity assay. Upon completion of the assay, we normalized the data to the H20S controls without the inhibitor to determine the impact of proteolytic activation with and without the inhibitor. Treatment with epoxomicin caused 95-100% inhibition of β5 in the absence and presence of all indicated 11S activators using the β5 substrate, LLVY-AMC (Fig 2A). However, PA28γ stimulation of the β2 substrate, RLR-AMC (T-L site) was essentially unaffected in the presence of epoxomicin. Alternatively, in the presence of the β2 inhibitor, leupeptin, our data demonstrated that β2 activity was diminished 94-100% in the absence or presence of all the indicated 11S activators, including PA28γ where activity went from 30-fold stimulation (Fig 2A) to no stimulation (Fig 2B). Our results definitively show that PA28γ’s ability to activate T-L activity does not require a functional CT-L site and cannot simulate T-L activity if the T-L like site is inhibited. Therefore, PA28γ does not function by altering the catalytic activity of β5 CT-L site as has been suggested for the PA200 activator.

**Figure 2.**
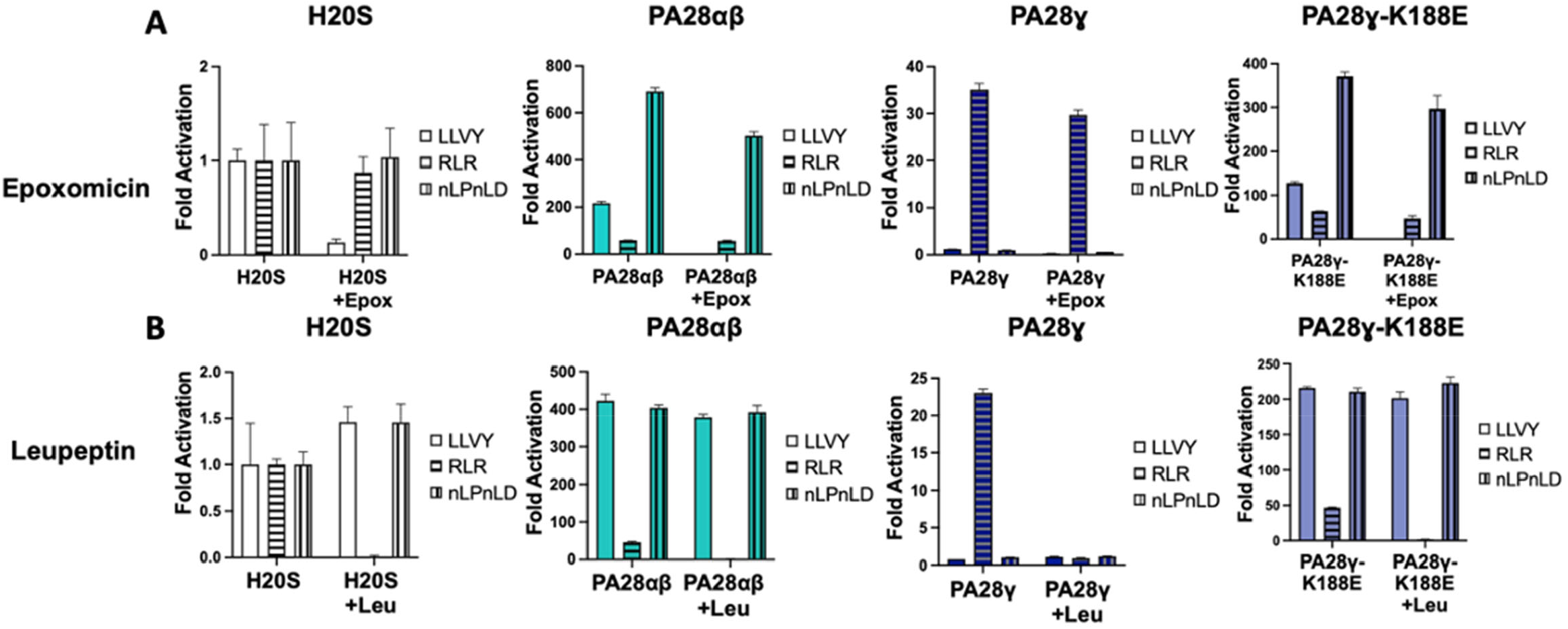
PA28γ does not change the specificity of other proteolytic sites to upregulate T-L activity. Purified H20S proteasomes were incubated with inhibitor [Epoxomicin (100nM) or Leupeptin (40μM)] and recombinant PA28αβ (50nM), PA28γ or PA28γ-K188E (62.5nM) were subsequently added to individual experiments. Proteasome activity was recorded for all proteolytic sites (RFU/min). Experiments were controlled for using buffer identical to the respective inhibitor. Results are the mean of at least three independent experiments performed in triplicate (error bars represent SD) normalized to the H20S control.

### PA28γ changes the K_m_ and V_max_ of T-L peptides for α3ΔN-Y20S

The results from our inhibitor data led us to ask the question of how PA28γ or PA28αβ changes proteolytic activity for each active site. To answer this question, we wanted to monitor the kinetic changes of each proteolytic site in the presence of either PA28γ or PA28αβ. Using the α3ΔN-Y20S, we were able to effectively answer this question as we can directly decouple the effects of activator binding and gate-opening from changes to the proteolytic sites. We performed dose responses of CT-L (LLVY-AMC), T-L (RLR-AMC), and C-L (nLPnLD-AMC) peptides to monitor kinetic changes of α3ΔN-Y20S proteolysis in the presence of either PA28γ or PA28αβ. Compared to the α3ΔN-Y20S controls, PA28αβ demonstrated negligible changes to the K_m_ or V_max_ for any of the three peptide substrates (Fig 3). This observation further demonstrates that PA28αβ does not allosterically alter the proteolytic sites, but functions solely as a gate-opening proteasomal activator. Alternatively, PA28γ does not modify the K_m_ or V_max_ for CT-L or C-L peptide substrates but does substantially change the K_m_ and V_max_ of the T-L substrate RLR-AMC. The presence of PA28γ decreases the K_m_ by >400% and increases the V_max_ by >300%. This equates to a >13fold increase in the catalytic efficiency (K_cat_/Km) of the T-L site by PA28γ. These results demonstrate that binding of PA28γ to the 20S increases both the catalytic affinity and the maximum proteolytic rate of the β2 (T-L) site, without affecting the Ct-L or C-L like activities. Therefore, PA28γ can use a non-gating activation mechanism, which is mechanistically distinct from PA28αβ. While it’s tempting to speculate that these results strongly suggest allosteric activation of the T-L site, similar results could possibly be obtained if PA28γ allowed the passage of some peptide substrates differently than others, to function as a substrate filter or sieve based on peptide charge. We will investigate this last alternative mechanism next.

**Figure 3.**
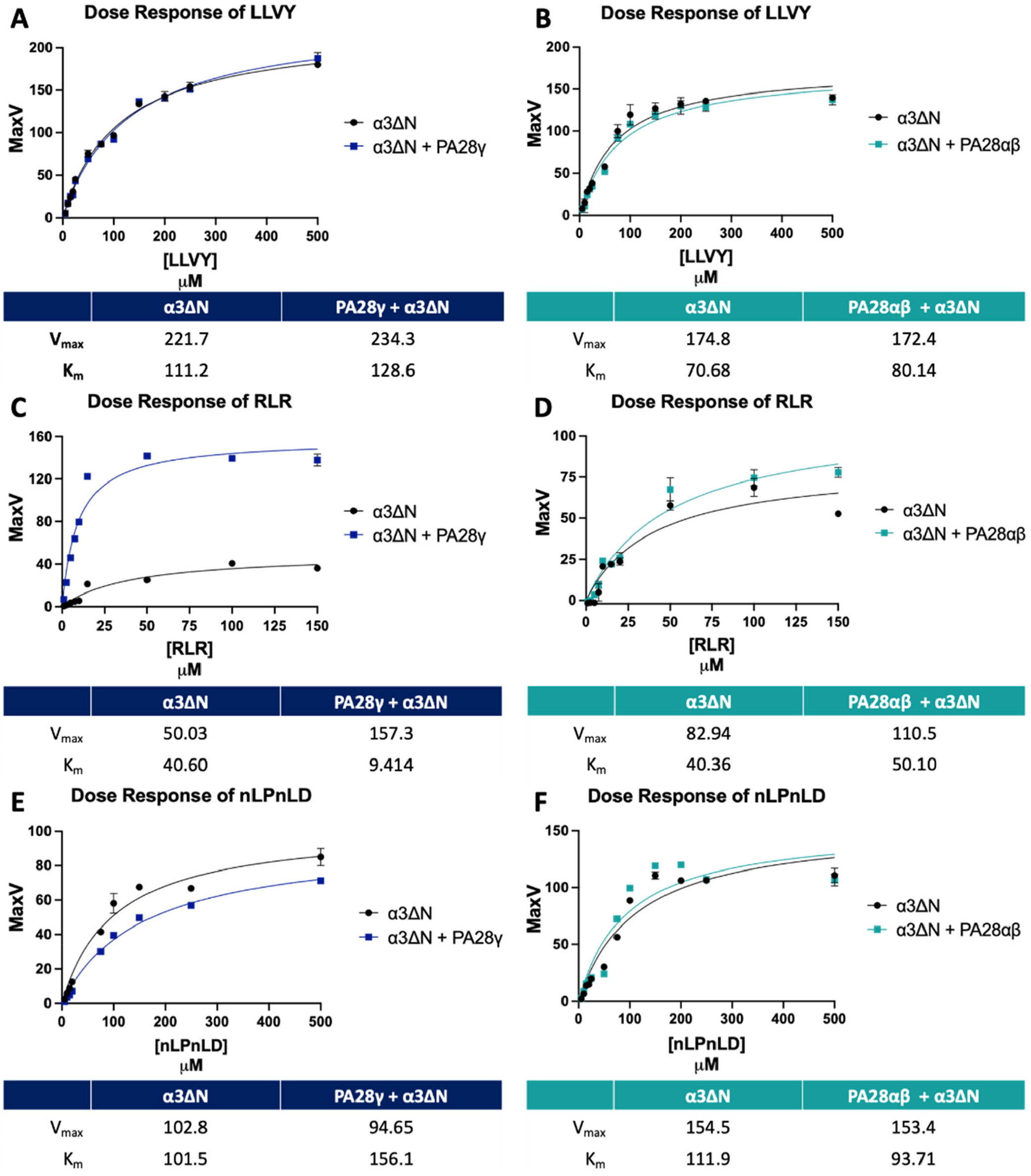
PA28γ changes the K_m_ and V_max_ of T-L peptides for α3ΔN-Y20S. Dose responses of peptide substrates were performed with α3ΔN-Y20S (0.1nM) and either PA28αβ (50nM) or PA28γ (62.5nM). Dose responses are as follows: A) PA28γ and LLVY-AMC (0-500μM), B) PA28αβ and LLVY-AMC (0-500μM), C) PA28γ and RLR-AMC (0-150μM), D) PA28αβ and RLR-AMC (0-150μM), E) PA28γ and nLPnLD-AMC (0-500μM), F) PA28αβ and nLPnLD-AMC (0-500μM). All experiments are the mean of three independent experiments (error bars represent SD).

### PA28γ does not select for substrates via its charged intrinsically disordered region or charged substrate entry pore

The above results indicate that PA28γ can upregulate T-L activity by a non-gating mechanism, but what remains unclear is whether PA28γ functions to exclude entry of some but not other substrates from the proteasome whereby, it could specifically allow entry of peptide substrates for the β2 active site^37^. All of the 11S/PA28 family members have an intrinsically disordered region (IDR), that has also been referred to as their homolog-specific insert, that is located around the substrate entry pore of the 11S complex and opposite of the proteasome-binding interface^37,38^. To answer if PA28γ can filter some peptide substrates, we designed two PA28γ variants that altered the IDR domain. The first replace the IDR of PA28γ with the IDR of PA28α, the second variant removed the entire IDR of PA28γ and replaced it with a small linker region sufficient to ensure formation of proper quaternary structure. We therefore created two PA28γ mutants, PA28γ-α and PA28γΔIDR. PA28γ-α is the PA28γ variant that contains the IDR from PA28α (i.e., it swaps IDR domains). A PA28α-γ swap was also created but did not heptamerize under our conditions and so could not be analyzed. The PA28γΔIDR is a deletion of PA28γ’s IDR but includes an 8-residue serine-glycine linker to limit the introduction of steric hinderance that a complete loss of the IDR would likely cause. When we tested the ability of these two PA28γ variants to activate 20S function we found that both the PA28γ-α and PA28γΔIDR upregulated β2 substrate degradation, similar to WT PA28γ, for both H20S and α3ΔN-Y20S. In addition, these 2 IDR variants showed no change in activity for the β1 or β5 substrates in the presence of either H20S or α3ΔN-Y20S, also similar to WT PA28γ (Fig 4 B&C). Based on these results we can conclude that PA28γ’s T-L activation mechanism does not require it’s IDR domain, and thus this domain must also not confer any peptide “filtering” capacity. In addition, the IDR domain from PA28α does not confer any gate opening activity to PA28γ when swapped.

**Figure 4.**
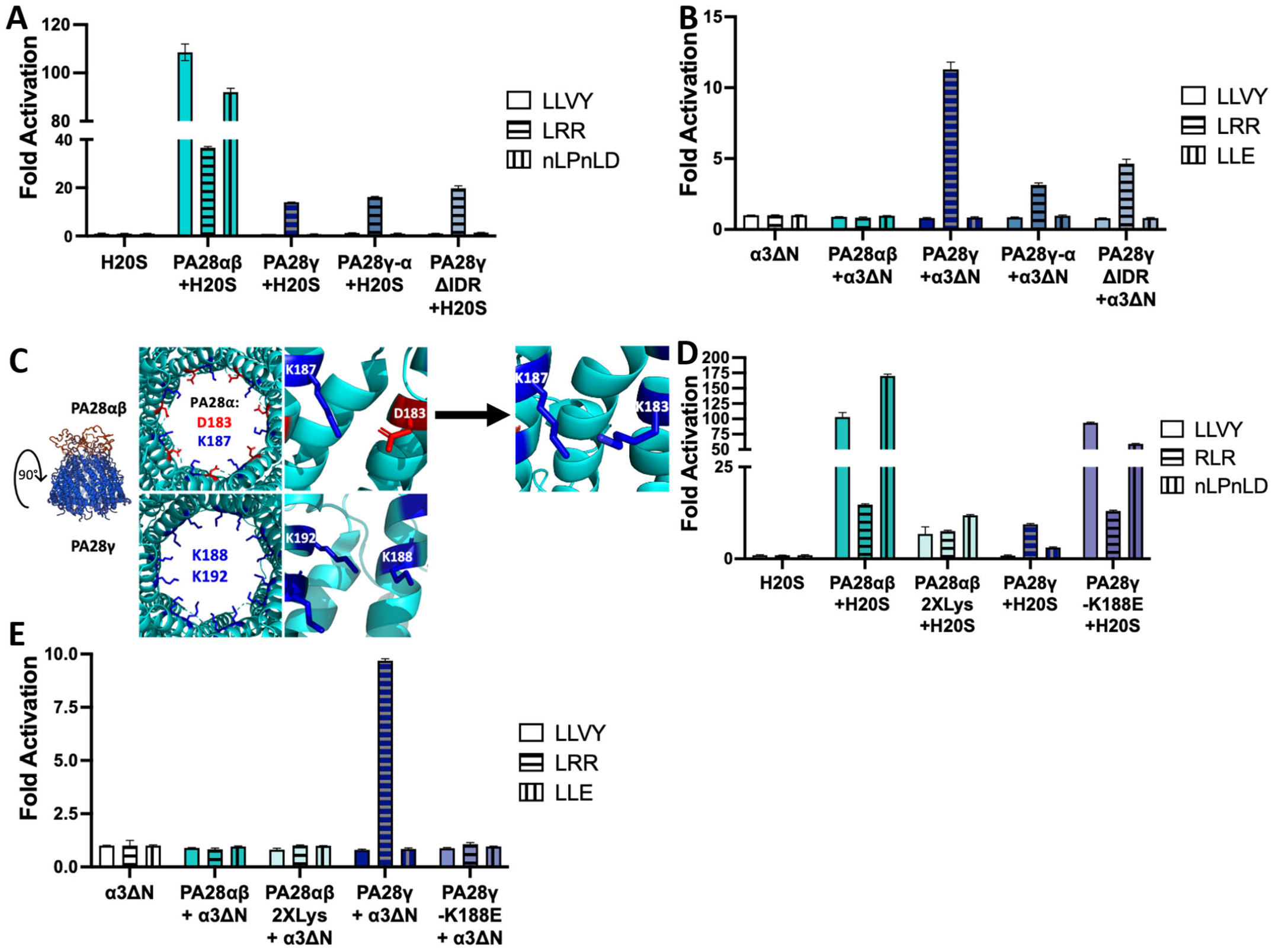
PA28γ does not select for substrates via its intrinsically disordered region or substrate entry pore. A) Purified H20S proteasome (1 nM) activity was measured for all three proteolytic sites (RFU/min) in the presence of recombinant PA28αβ (50nM), PA28γ, PA28γ-α, or PA28γΔIDR (62.5nM). B) Purified constitutively open-gate α3ΔN-Y20S (0.1nM) activity was measured for all three proteolytic sites (RFU/min) in the presence of recombinant PA28αβ (50nM), PA28γ, PA28γ-α, or PA28γΔIDR (62.5nM). C) Site directed mutagenesis was used to create the PA28αβ mutant, PA28αβ2XLys, a substrate entry pore mutant that maintains positive charges around the ring, like PA28γ. (PDB: 5MX5; PA28γ structures based on modified PDB: 5MX5) D) Purified H20S proteasome (1nM) activity was measured for all three proteolytic sites (RFU/min) in the presence of recombinant PA28αβ, PA28αβ2XLys (50nM), PA28γ, or PA28γ-K188E (62.5nM). E) Purified constitutively open-gate α3ΔN-Y20S (0.1nM) activity was measured for all three proteolytic sites (RFU/min) in the presence of recombinant PA28αβ, PA28αβ2XLys (50nM), PA28γ, or PA28γ-K188E (62.5nM). All experiments are the mean of three independent experiments (error bars represent SD).

As discussed previously, PA28γ-K188E has a substrate entry pore that mimics the substrate entry pore of PA28αβ and changes the functional mechanism of PA28γ from allosteric activation to gate-opening (Fig. 1A). We therefore asked the question if the charges in this pore region could affect which types of peptides are able to pass through PA28αβ. To ask this question we introduced point mutations into PA28αβ that would mimic the charges in the substrate entry pore of PA28γ. We then asked if these mutations would be sufficient to switch PA28αβ’s activity to that of PA28γ. This experiment was performed for two reasons: 1) to test whether PA28αβ and PA28γ can filter peptide substrates through their substrate entry pores, and 2) to implicate helix 3 in the unique functional activity of PA28αβ and PA28γ on the 20S proteasome. As PA28αβ is two separate subunits, we created this mutant by inducing point mutations in helix 3 of PA28α-D183K and PA28β-D173K, which will further be called PA28αβ2XLys (Fig 4C). A proteasome activity assay using WT H20S revealed that PA28αβ2XLys still stimulates the degradation of all three types of substrates though at a reduced capacity: approximately 2-10-fold less activity amongst all peptide substrates when compared to WT PA28αβ (Fig 4D). Nevertheless, these results demonstrate that PA28αβ2XLys is functional to induce gate opening, but importantly does not selectively prevent the passage of some types of peptides, despite its very different pore charges. To confirm this conclusion we also found that, PA28αβ2XLys was unable to stimulate peptide hydrolysis in the α3ΔN-Y20S at any capacity (similar to WT PA28αβ), which also indicates that PA28αβ2XLys maintains its gate opening ability (Fig 4E) but does not “filter” substrates based on their charge. This result reveals that that the generation of a completely positively charged pore in PA28αβ was not able to switch it to a T-L activating complex (like PA28γ), thus demonstrating that differences in pore charge are unable to selectively exclude some types of peptides from entering the degradation chamber. One caveat of this experiment is that mutagenesis of helix 3 in PA28αβ diminished the activator’s ability to induce gate-opening, but it did not change its general function. Based on the structure of PA28γ, there are 3 different regions where peptide passage through PA28γ and entry into the 20S could be affected: 1) the IDR, 2) the substrate entry pore, and 3) the proteasome gate. We have generated mutants in all three of these locations and found that none of them could change the effect that PA28γ has on the proteasome. Therefore, these results clearly demonstrate that PA28γ does not select for peptide substrates through its IDR or its substrate entry pore and does not specifically open the proteasomal gate for T-L peptide substrates, but, in fact, allosterically upregulates the catalytic capacity of the β2 subunits thus increasing T-L-specific peptide proteolysis.

### Cryo-EM electron density reveals structural topology of PA28γ

Recently, the structures of various 11S family members have been solved using Cryo-EM and X-Ray Crystallography^13,39,40^. However, the structure of the PA28γ complex or the PA28γ-20S proteasome complex has remained undetermined. Using Cryo-EM, we were able to generate a low-resolution reconstruction of the human PA28γ-20S complex to 4.3Å (Fig 5A). This electron density reveals that PA28γ binds to the 20S proteasome using its C-termini to dock into the proteasomal intersubunit pockets of the α-ring, like PA26^41^ and PA28^13,40^ activators (Supplemental Figure 2). Further, the recent structure of the PA28αβ-immunoproteasome complex fits fairly well into our PA28γ-20S structure map showing that PA28γ adopts an overall structure that is similar to PA28αβ (Fig 5C). Further, when we overlay our electron density with the recent structure of the PA28αβ-immunoproteasome complex, it is clear that PA28γ adopts an overall topology and quaternary structure that is similar to PA28αβ (Fig 5C). The tertiary and secondary structures of PA28αβ and PA28γ also align similarly relative to one another (Fig 5D and E). Figure 5E also shows a slice through the and PA28γ map that includes the PA28αβ model, and it’s clear the helix 3 (center most helix) occupies similar space in both homologs. Interestingly, our structure reveals a density in the pore of PA28γ, that does not appear in the model of PA28αβ (Fig 5B and D). This Cryo-EM structural reveals that PA28γ adopts a similar tertiary conformation to the fellow mammalian homolog, PA28αβ, even though PA28γ only has 42.1% sequence identity to PA28α and 33.6% identity with PA28β, and even though both activators activate proteasome function in very different ways.

**Figure 5.**
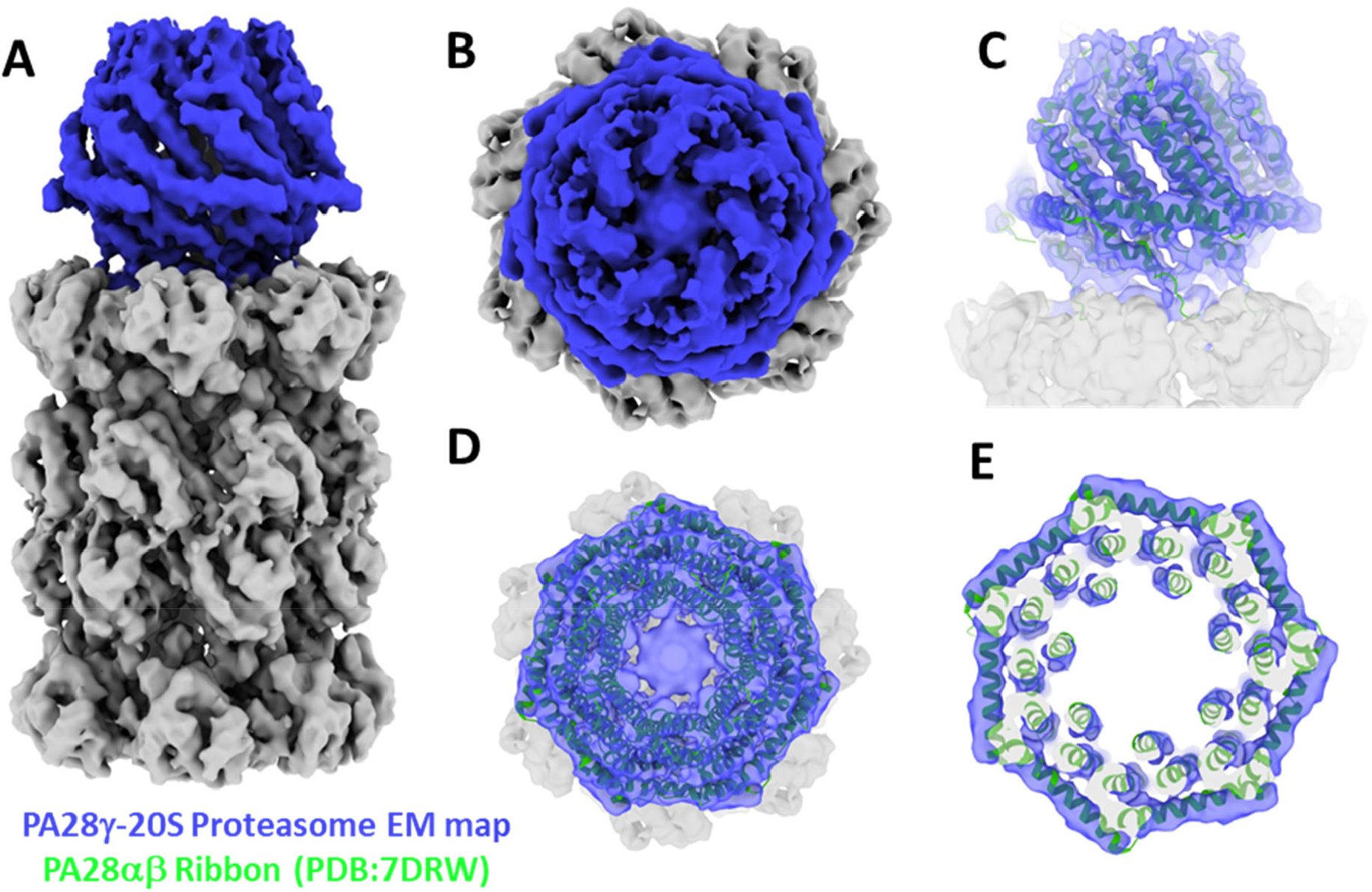
Cryo-EM reveals PA28γ has a similar structure to other 11S regulators when bound to the 20S proteasome. A) Full 4.3Å electron density of the human PA28γ-20S proteasome complex after density modification. PA28γ density is colored blue, and the 20S densities are colored grey. C7 symmetry was applied during 3D reconstruction since PA28γ is a homoheptamer. B) Top-down view of the substrate entry pore of PA28γ. C) Molecular model of PA28αβ (PDB: 7DRW) fit into the electron density of the PA28γ-20S complex (side view). D) Top view of C. E) Same as D except the map and model are cropped through the 7-fold symmetry axis to focus on the Helix 3 density (center most helix).

## Discussion

This study unambiguously demonstrates that within the 11S family, PA28γ and PA28αβ can use two completely different mechanisms to activate proteasome function. Previous work initially determined that PA28γ could stimulate the degradation of trypsin-like substrates but did not explicitly define how it achieves its function. Based on previous literature and structure/function analysis, five potential mechanisms were proposed that could answer how PA28γ upregulated T-L peptide degradation: 1) specific substrate selection, 2) allosteric upregulation of the T-L catalytic site, 3) allosteric inhibition of the CT-L and C-L catalytic sites, 4) conversion of the CT-L or C-L sites to new T-L catalytic sites, and 5) gate-opening in combination with a previous hypothesis. Our results effectively ruled out mechanism 1 that the PA28γ could function as a substrate selective “sieve” and allow only entry of only peptide substrates with positive charges. The “sieve” model would require that PA28γ is able to open the 20S gate, but only allow entry of select peptides (e.g., T-L peptides). Further, if PA28γ functioned as a “sieve” it would also limit CT-L and C-L peptides from entering the proteasome, which would appear as inhibition in the open channel 20S. Our results from Figure 1D clearly demonstrate that PA28γ upregulates the degradation of the T-L peptide but did not inhibit the degradation of CT-L or C-L peptides in the constitutively active proteasome. Moreover, it is reasonable to assume that if PA28γ was using a sieve model, that the IDR surrounding the substrate entry pore may play a role in substrate selection. Figure 4 clearly shows that mutagenesis to the IDR does not affect PA28γ’s ability to upregulate the proteolysis of T-L peptide substrates. In addition, mutation of PA28αβ’s pore to generate a ring of positive lysine’s mimicking PA28γ’s pore (Fig 4C-E) demonstrated that this 2XLys mutation did not confer any “sieving-like” properties to PA28αβ and did not change how it activated 20S function. These results clearly rule out contributions from mechanism 1 for PA28γ function. None of our experiments could rule out mechanism 2—allosteric activation of the T-L like site. Mechanism 3 or allosterically inhibition of CT-L or C-L activity (combine with gate-opening—mechanism 5) to increase T-L degradation is ruled out by Figure 1 showing that T-L activity is still stimulated even when the gate is constitutively open and CT-L and C-L activity are not affected in the open mutant. In addition, Figure 3 (A & E) show that the equilibrium kinetics of CT-L and C-L peptides remain unchanged when PA28γ is bound to open channel proteasome complex demonstrating that PA28γ cannot inhibit these active sites (Fig 3C serves as a control to show that PA28γ does indeed bind to the open channel 20S in these conditions). Together figure 3 conclusively demonstrated that in the absence of a functioning 20S gate, PA28γ increased the catalytic affinity (K_m_) and increased the V_max_ of the T-L like site for RLR-AMC (Fig 3C), inducing a 13-fold increase in catalytic efficiency of the T-L site. These results suggest that PA28γ can allosterically activate the T-L proteolytic site. Mechanism-4 is ruled out by results shown in figure 2 that shows that PA28γ does not switch the CT-L site to be more T-L. These experiments demonstrate that PA28γ’s ability to upregulate T-L peptide degradation was unaffected even when the CT-L site was modified with a covalent inhibitor, revealing that PA28γ could not switch the CT-L site to T-l like, as has been suggested for PA200’s mechanism of T-L like activation. In addition, we further verified that pretreatment with a T-L inhibitor could indeed inhibit PA28γ’s ability to upregulate T-L activity. Finally, the fact that PA28γ’s ability to upregulate T-L peptide degradation occurs with WT or open channel mutant 20S (Figure 1C & D) demonstrates that it does not need to induce gate opening to stimulate T-L activity (ruling out mechanism 5), and therefore it must do so via allosterically altering the proteolytic site activity. Taken together these results clearly demonstrate that PA28γ, through long-range interactions (Fig 6A), must allosterically affects the β2 subunits of the 20S proteasome to upregulate its trypsin-like activity.

**Figure 6.**
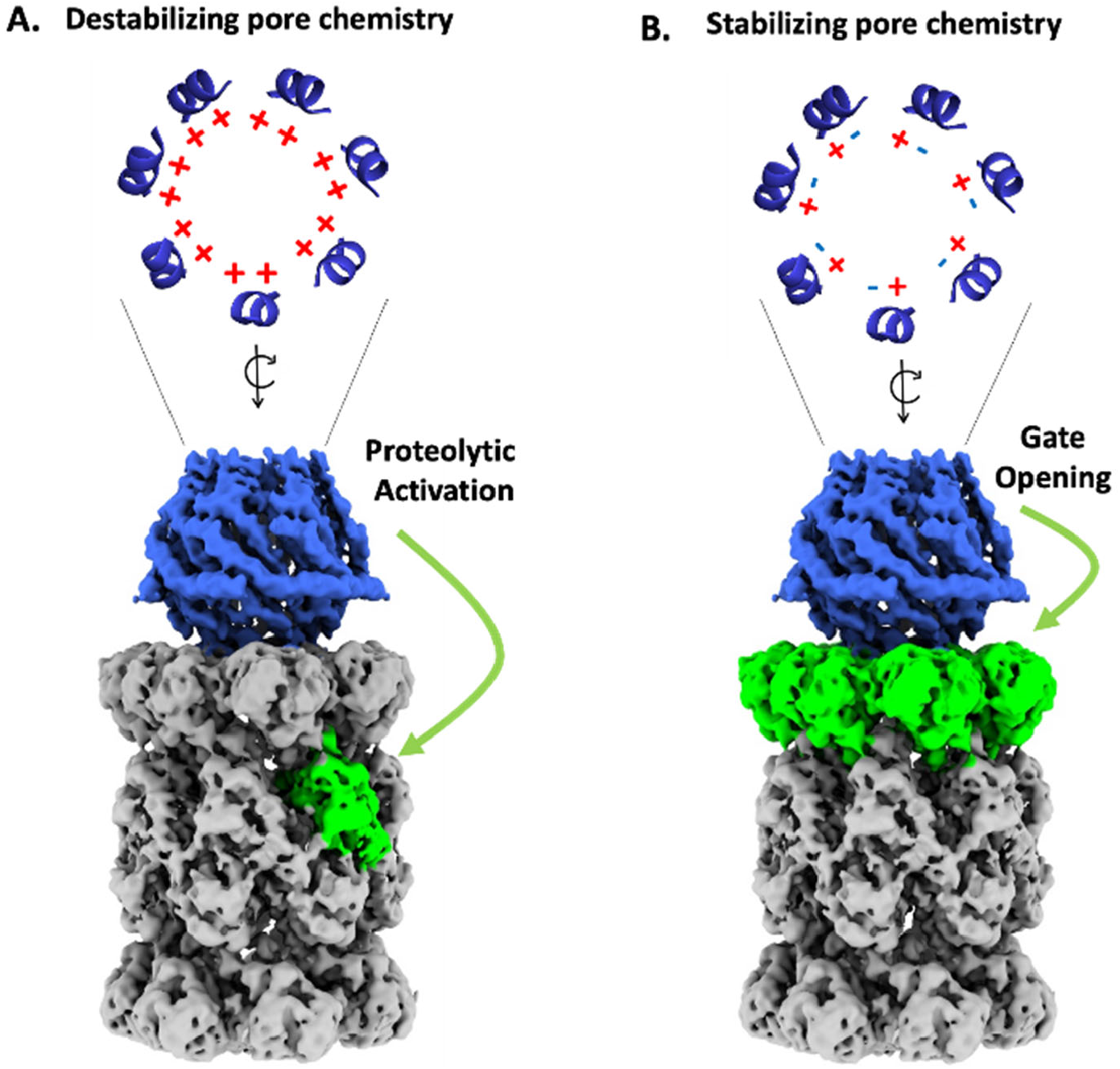
Model of PA28γ and PA28γ-K188E pore charges and their effects on the 20S proteasome. Proteasomal regulation by PA28γ is directly affected by helix-3 interactions. A) WT PA28γ has a positively charged (i.e., helix-3 repelling), substrate entry pore which allosterically induces proteolytic activation of the T-L proteolytic site in the 20S proteasome. B) A single point mutation (K188E) to PA28γ creates an ionically stabilized (helix-3 interacting) substrate entry pore that alternates positive and negative charges similar to PA28αβ. Replacing these charges with hydrophobic residues functions similarly. This changes the regulatory mechanism of PA28γ causing to switch its function to one that induce proteasomal gate opening. These conclusions are based on PA28γ’s ability to affect the activity a the open-channel 20S proteasome, which can dissociate gating affects from active site affects.

WT PA28γ has two basic residues on helix 3 (K188 and K192) while WT PA28αβ has a basic (lysine) and acidic (aspartate) residue in the homologous positions of PA28α (D183 and K187) and PA28β (D173 and K177; Fig 1, Fig 6). The structures of PA28αβ reveal that the basic residue in helix 3 interacts with the acidic residue in helix 3 of the neighboring subunit^15^, which would effectively induce a repeating salt-bridge around the substrate entry pore that could theoretically stabilize the substrate entry pore and the 11S structure in general. We hypothesize that the rigidity within the substrate entry pore, due to stabilizing salt bridges, could somewhat lock PA28αβ (and other similar gate-opening 11S regulators) into a conformation that would allow for the activation loops to more effectively impose their symmetry onto the α-subunits of the 20S to cause gate-opening as has been proposed for PA26^40^. Here we clearly show that PA28αβ’s only function is to induce gate opening since it could not affect open channel 20S activity. Contrarily, PA28γ’s helix 3 provides repeating positive charges that surround the substrate entry pore, which is expected to generate repulsive forces that reduce pore stability and create a more dynamic or flexible 11S activator. In support of this notion the K188E mutation in PA28γ introduces stabilizing interactions to the Helix 3 ring, which switches PA28γ’s function to gate-opening (Fig 1, 4 & 6). We further probed these helix 3 interactions and switching mechanism by generating the PA28γ2XCys mutant, which contains two hydrophobics in the K188 and K192 positions. This variant similarly switches PA28γ from T-L activating to gate-opening (whether disulfide crosslinked or not). Therefore, two different mutations that increase the interactions of helix 3 with neighboring helix 3’s, likely increase helix 3 positional stability in the pore, and this causes PA28γ to switch into a gate-opening state. In contrast, when we destabilized the substrate entry pore of PA28αβ with our PA28αβ2XLys mutant, we saw a marked reduction in gate-opening activity (mirroring PA28γ activity) when compared to WT PA28αβ, but it did not induce a complete functional switch like the PA28γ-K188E mutant did, meaning it did not induce T-L activity (Fig 4D). This indicates that there must be other differences between PA28αβ and PA28γ that compensate for their mechanistic differences besides pore stability. Although mutations that increase helix 3 interactions in the pore of PA28γ can cause it to switch activity, it is not known if WT PA28γ, under physiological conditions, can similarly switch states as part of its normal function to degrade proteins, though it is tempting to speculate that this switching mechanism could be a component of how it regulates protein degradation, which is currently not understood.

Based on our observations, we hypothesize that PA28γ’s positively charged substrate entry pore allows it to adopt a conformational that can generate the long-range allosteric effects that allow it to activate the 20S T-L site. In support of this notion, a recent study by Lesne et. al used Hydrogen-Deuterium eXchange coupled to Mass Spectrometry (HDX-MS) to determine that PA28αβ and PA28γ adopt different conformations when binding the proteasome, as the activation loops and helix 1 of PA28αβ were heavily protected from HDX solvent exchange when bound to the proteasome compared to PA28γ^42^. Our structure of the human PA28γ-20S complex reveal that PA28γ adopts a quaternary topology similar to other structurally characterized 11S regulators (i.e., PA26, PA28*Pf*, and PA28αβ). Further, the overall structure of the PA28γ-20S proteasome complex revealed that PA28γ docks its C-termini into the intersubunit pockets of the proteasome’s α-ring, a mechanism paralleled by other 11S regulators, and does not bind the β ring directly to upregulate T-L proteolysis. We also revealed a density unique to PA28γ that occupies the substrate entry pore. It is expected that this density is its IDR, and although we could not resolve this region, it supports the idea that this region is highly disordered. Our structure allows us to visually conclude that PA28γ in general has many overall qualities that align with the 11S family of regulators. While the application of C7 symmetry to the PA28γ complex seems reasonable since PA28γ is a homoheptamer, the same symmetry application to the heteroheptomeric 20S obviously causes loss of any meaningful asymmetric information for the 20S, which would be required to observe T-L structural changes. Unfortunately, the particle number obtained was too low for 3D reconstruction without symmetry application. Lesne et. al’s HDX-MS study complements our biochemical results and supports our hypothesis that PA28γ uses a distinct proteasomal activation mechanism from other 11S regulators, while structurally preserving similar tertiary and quaternary protein structures.

We set out to definitively determine what mechanism PA28γ uses to regulate peptide degradation through the 20S proteasome. The results presented here explicitly demonstrate that PA28γ can allosterically and selectively activate the 20S proteasomes T-L proteolytic site, without affecting the CT-L or C-L proteolytic sites (Fig 1A&B, Fig 3). Further, these data show that PA28γ’s helix 3 plays a major role in regulating the overall function of the activator (Fig 1, Fig 4, and Fig 5). In addition, PA28γ-20S complex plays a role in nuclear processes such as: cell cycle progression and cell proliferation^43,44^, apoptosis^43^, formation of nuclear speckles^45^, and DNA repair^46^. Within these processes, PA28γ-20S complex has also been demonstrated to facilitate the degradation of many nuclear proteins, such as SRC-3/AIB1^47^, c-Myc^48^, p21^49^, Hepatitis C Virus (HCV)^50^, and MDM2^51^. Interestingly, many PA28γ substrates appear to be DNA binding proteins. In addition to cancer progression, PA28γ could also play a role in protecting against some neurodegenerative diseases, like Huntington’s disease, which is caused by expression and accumulation of the PolyQ expanded Huntington protein^52^. A prior study using PolyQ expanded peptide substrates determined that the T-L site was responsible for PolyQ substrate degradation, and longer PolyQ expanded repeats (30 Q amino acids) were unable to be degraded by the mammalian proteasome^32^. While further studies showed that the proteasome could cleave PolyQ proteins^53^, it’s clear from both studies that PolyQ cleavage is very slow. Based on these results, it seems that PA28γ’s ability to stimulate the T-L site would be expected to play an important role in accelerating the degradation of poly-Q proteins, which could be protective. In support of this notion, treatment with PA28γ has been shown to have a protective effect in PolyQ neurodegenerative disease studies^7,8^. Therefore, understanding how PA28γ enhances the proteolytic activity of the T-L site as shown here would provide a mechanistic understanding and platform to design drugs that could be used to potentially treat Huntington’s disease. Therefore, based on our results and this literature, we hypothesize that the role of PA28γ in proteasomal nuclear proteostasis is to enhance the T-L activity of the 20S proteasome for more effective degradation of positively charged proteins, which are often DNA and RNA binding proteins. This proposed function of the PA28γ-20S complex could explain why some cancers upregulate PA28γ, as it could facilitate the expeditious degradation of important transcription factors and ubiquitination cascade proteins. This function could also answer the question of why PA28γ overexpression in PolyQ neurodegenerative disease rescues the disease phenotype. This study lays the groundwork to better understand the mechanism of how PA28γ activates T-L activity, which could inform efforts to design inhibitors of PA28γ function that could be used to specifically treat PA28γ-overexpressed cancers, and 20S proteasome T-L site stimulating drugs that could be used to treat PolyQ neurodegenerative diseases.

## Methods

### Proteasome purifications

<u>Bovine 20S (B20S)</u> proteasomes were purified from bovine liver as described^54^. Liver was homogenized, cleared, and passed over weak anion exchange resin (DE52, GE Life Sciences). B20S proteasomes were eluted using a stepwise NaCl gradient. Fractions with proteolytic activity were pooled and dialyzed before strong anion exchange (Resource Q, GE Life Sciences) separation using a linear NaCl gradient. Elution with significant suc-LLVY-AMC activity were pooled and further separated using a hydroxyapatite chromatography column (CHT-I, Bio-Rad) using a linear KPO_4_ gradient. Eluted fractions with significant proteolytic activity were pooled, dialyzed, and B20S purity (>95%) was determined using SDS-PAGE and densitometry using ImageJ. <u>Eukaryotic α3ΔN-Y20S</u> were expressed and purified from yeast using anion exchange chromatography, as described with minor modifications^55^ Briefly, size-exchange chromatography (SEC) (Superose 6 Increase, GE Life Sciences) was performed on pooled fractions with proteolytic activity after Resource Q. <u>Human 20S (H20S)</u> proteasomes were purified from stably transfected HEK 293T cells, as previously described^56^.

### Proteasome activator purifications

<u>Proteasome activator (PA) 28αβ</u> was purified as described^57^. Briefly, recombinant PA28αβ was expressed in BL21-STAR *E*. coli and purified using strong anion exchange (HiTrapQ and MonoQ, GE Life Sciences), followed by hydroxyapatite (CHT-II, Bio-Rad), and finished with SEC (Superose 6 Increase, GE Life Sciences). <u>Recombinant PA28γ</u> was expressed in Rosetta *E. coli* and purified using Ni-NTA affinity resin (Quiagen) and followed with SEC (Superose 6 Increase, GE Life Sciences), as previously described^10^. PA28γ-K188E was created using QuikChangeII Site Directed Mutagenesis Kit (Aligent) and purified using methods like PA28γ, as previously described^30^. <u>PA28γ-α, PA28γ-2XCys, PA28αβ2XLys, and PA28γΔIDR</u> constructs were designed as G-Blocks with N-terminal 6XHis Tags (Integrated DNA Technologies) and cloned into pET11a plasmids. pET11a plasmids with successfully cloned G-blocks were transformed into BL21-STAR *E. coli* and purified using the PA28γ Ni-NTA purification methods.

### Proteasome activity assays

Unless otherwise stated, B20S (1 nM), Y20S (1 nM), H20S (1 nM), and α3ΔN-Y20S (1 nM) were all assayed using fluorogenic peptides, as previously described^36^, using a Biotek 96-well plate reader. Briefly, 20S proteasomes were incubated in a reaction buffer containing 50 mM Tris-HCL (pH 7.5), 5% glycerol, 1 mM DTT, and 100 μM fluorogenic substrate (suc-LLVY-AMC, boc-RLR-AMC, boc-LRR-AMC, Ac-nLPnLD-AMC or Z-LLE-AMC) and put into a 96-half well black flat-bottomed treated plate. 20S proteasomes were subsequently treated with either PA28αβ (50 nM), PA28γ (62.5 nM), PA28γ-K188E (62.5 nM), PA28αβ2XLys (50nM), PA28γ-α (62.5 nM), or PA28γΔIDR (62.5nM). Fluorescence measurements were taken every 30s for 60 minutes (ex/em: 380/460 nm). The slope of the linear increase in fluorescence is directly proportional to the rate of 20S proteasome activity. Assays in the presence of proteasomal activators are normalized to the 20s proteasome only control. All molar concentrations of the proteins above are based on the molecular weight of the total complex.

### Proteasome inhibitor assays

Assays were performed under reaction conditions and protein concentrations similar to proteasome activity assays. B20S proteasomes were incubated in reaction buffer with either PA28αβ, PA28γ, or PA28γ-K188E and either Epoximicin (100 nM) or Leupeptin (40μM) was subsequently added (Epoximicin, Enzo Life Sciences; Leupeptin, Sigma Aldrich). Assays were controlled for using replicates without inhibitors read simultaneously.

### Substrate dose responses

Dose response assays were done following a modified proteasome activity assay protocol. Briefly, α3ΔN-Y20S (1 nM) was put into a proteasome activity assay buffer alone or with either PA28αβ (50nM) or PA28γ (62.5nM). Fluorogenic substrate was added in a dose dependent manner (suc-LLVY-AMC: 0-500μM; boc-RLR-AMC: 0-200 or 225μM; ac-nLPnLD-AMC: 0-500μM). Assays were analyzed for the MaxV at each concentration and dose responses were analyzed using GraphPad Prism9.

### Oxidation/Reduction assay

WT PA28γ and PA28γ2XCys were assayed using a modified protocol^35^. Briefly, WT PA28γ and PA28γ2XCys were incubated in 0.1% β-Mercaptoethanol for 1 hour before being desalted using a Zeba™ Spin Desalting Column (Thermo Scientific). Proteins were then incubated at 37°C for 10 minutes in 0.1% β-Mercaptoethanol or Tetrathionate (1mM) to reduce or oxidize the cysteines, respectively. After incubation, proteins were added to a proteasome activity assay master mix previous protocol was followed. Further, reduced, or oxidized proteins were run on a 4-12% Bis-Tris Gradient SDS gel, and subsequently stained for total protein (Colloidal Blue) or transferred and blotted using western blot methods. Proteins were transferred at room temperature to a PVDF membrane using 1X NuPAGE Transfer Buffer. Upon transfer, membranes were blocked with 10% milk, and then probed with α-PSME3 antibody (Bethyl Laboratories) overnight. Membrane was washed and further probed with an HRP 2° antibody and imaged using the Syngene GeneSys B:Box imaging system and software.

### Cryo-EM Sample Preparation and Data Collection

Copper Quantifoil R 1.2/1.3 300 mesh (EMS) grids were cleaned and treated with amylamine using a PELCO easiGlow Glow Discharge cleaning system. PA28γ and H20S were mixed at a 1:1 molar ratio and 3 uL of the sample mixture was placed onto a grid. Grid was subsequently blotted by hand using blotting paper, and immediately flash frozen in liquid ethane using a manual plunge freeze apparatus. Data collection was done using a Titan Krios transmission electron microscope (Thermo Fisher) operated at 300kW and a magnification of x81,000, which resulted in 1.08Å/px. Images were collected using a Falcon IIIEC direct electron detector camera equipped with a K3/GIF system operating in counting and super resolution modes. Electron dose per pixel of 50 e-/Å^2^ was saved as a movie with each movie being dose-fractioned into 40 frames within a target defocus range of -2.5 to -1.25. All data was collected using cryoSPARC software (Thermo Fisher).

### Cryo-EM Single Particle Analysis

2328 total movies were collected, and we used 2300 for structural determination. Single particle analysis of the PA28γ-20S proteasome complex was done using cryoSPARC (Supplemental Figure S3). All images were aligned and summed using motion correction. After CTF estimation using cryoSPARC’s patch-based CTF estimator, 31712 particles were autopicked from the micrographs and subjected to several rounds of 2D classification to get rid of junk particles (Supplemental Figure S3B). 3358 particles primarily side view orientation (Supplemental Figure S3E) from single and double cap in PA28y-20S 2D classes were used to generate five ab-initio models to further remove junk particles (Supplemental Figure 3C). One of the resulting ab-initio models produced a PA28y-20S complex from 876 particles, which were used for homogeneous regiment for 3D classification with C7 symmetry applied using the map from prior determined ab-into model (Supplemental Figure 3C &D). The gold standard (0.143) FSC resolution was calculated form 2 half maps (Cryosparc) to be 4.4 Å (Supplemental Figure 3F). All representations of the PA28γ-20S proteasome complex were created using UCSF ChimeraX^58^

## Acknowledgements

Transmission electron micrographs were recorded at the University of Virginia Molecular Electron Microscopy Core facility (RRID:SCR_019031), which is supported in part by the School of Medicine and built with NIH grant G20-RR31199. In addition, the Titan Krios (S10-RR025067), Falcon II/3EC direct detector (S10-OD018149), and K3/GIF (U24-GM116790) were purchased in part or in full with the designated NIH grants. Molecular graphics and analyses performed with UCSF ChimeraX, developed by the Resource for Biocomputing, Visualization, and Informatics at the University of California, San Francisco, with support from National Institutes of Health R01-GM129325 and the Office of Cyber Infrastructure and Computational Biology, National Institute of Allergy and Infectious Diseases. This work was supported by NIH-R01GM107129 to D.M.S.

## Supplement

**Figure S1.**
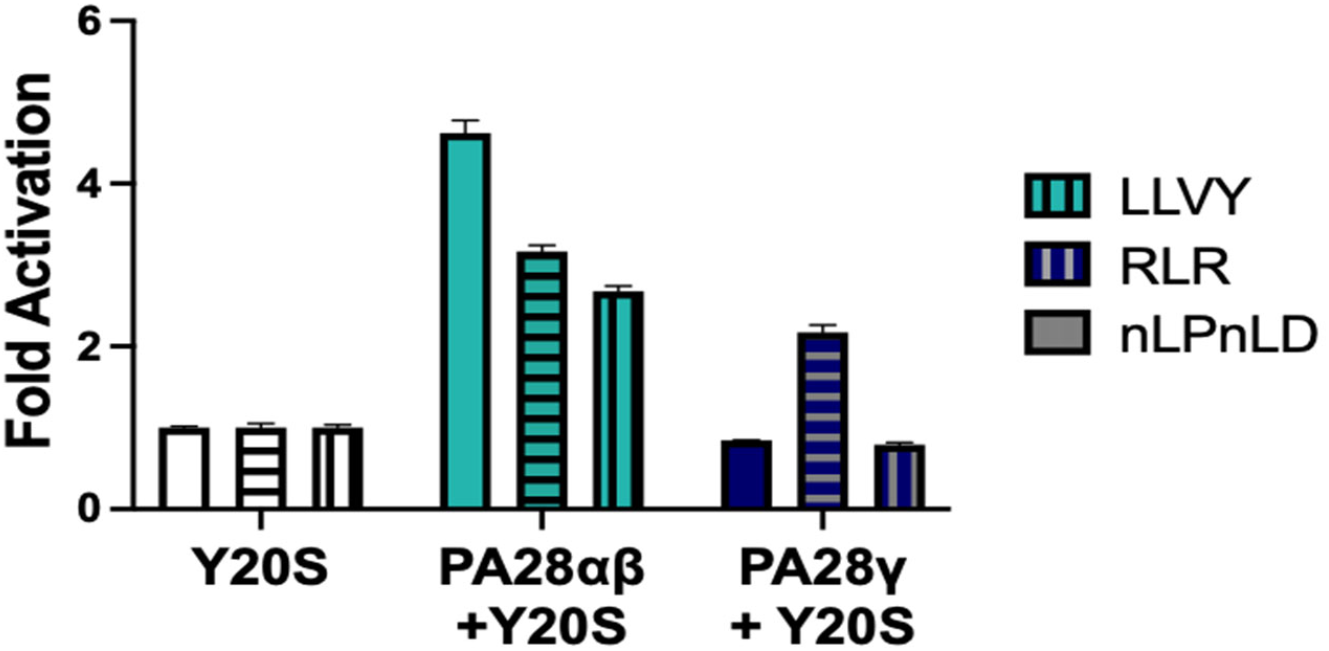
WT Y20S Proteasome Activity Assay. Proteasome activity assay with PA28αβ and PA28γ reveals a similar pattern of activation between Y20S and H20S (Fig 1C). Purified Y20S (1nM) was tested for all three proteolytic sites (RFU/min) in the presence of recombinant PA28αβ (50nM), and PA28γ (62.5nM). Results are the mean of at least three independent experiments performed in triplicate (error bars represent SD) normalized to the Y20S control.

**Figure S2.**
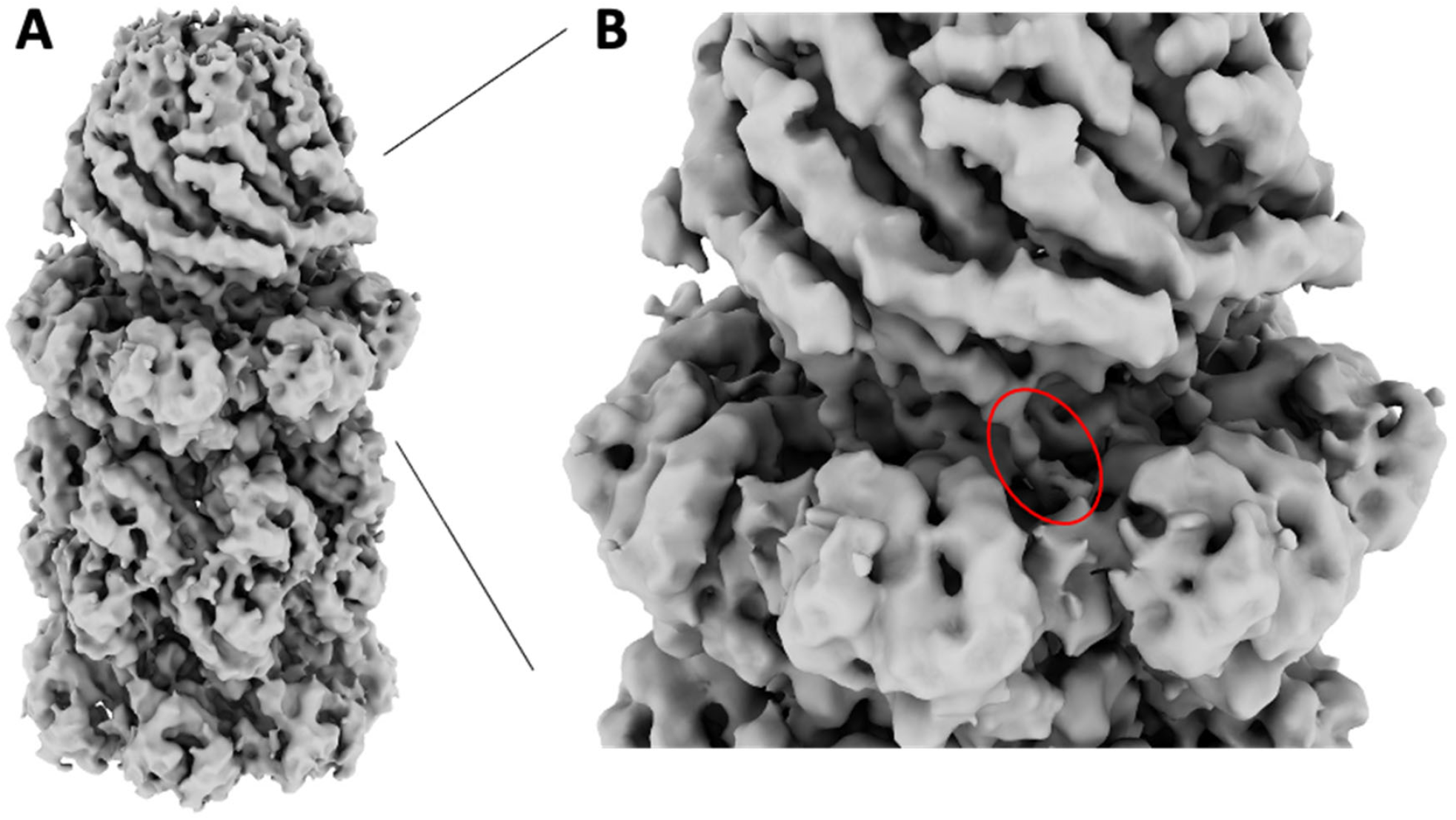
Cryo-EM Map and C-terminal Docking Site. PA28γ-20S complex shows C-termini are docked into the intersubunit pockets of the 20S. A) Complete side view, B) zoomed view of “A” shown by lines. Red circle highlights one of PA28γ’s 7 C-termini docking into one 20S intersubunit pocket.

**Figure S3.**
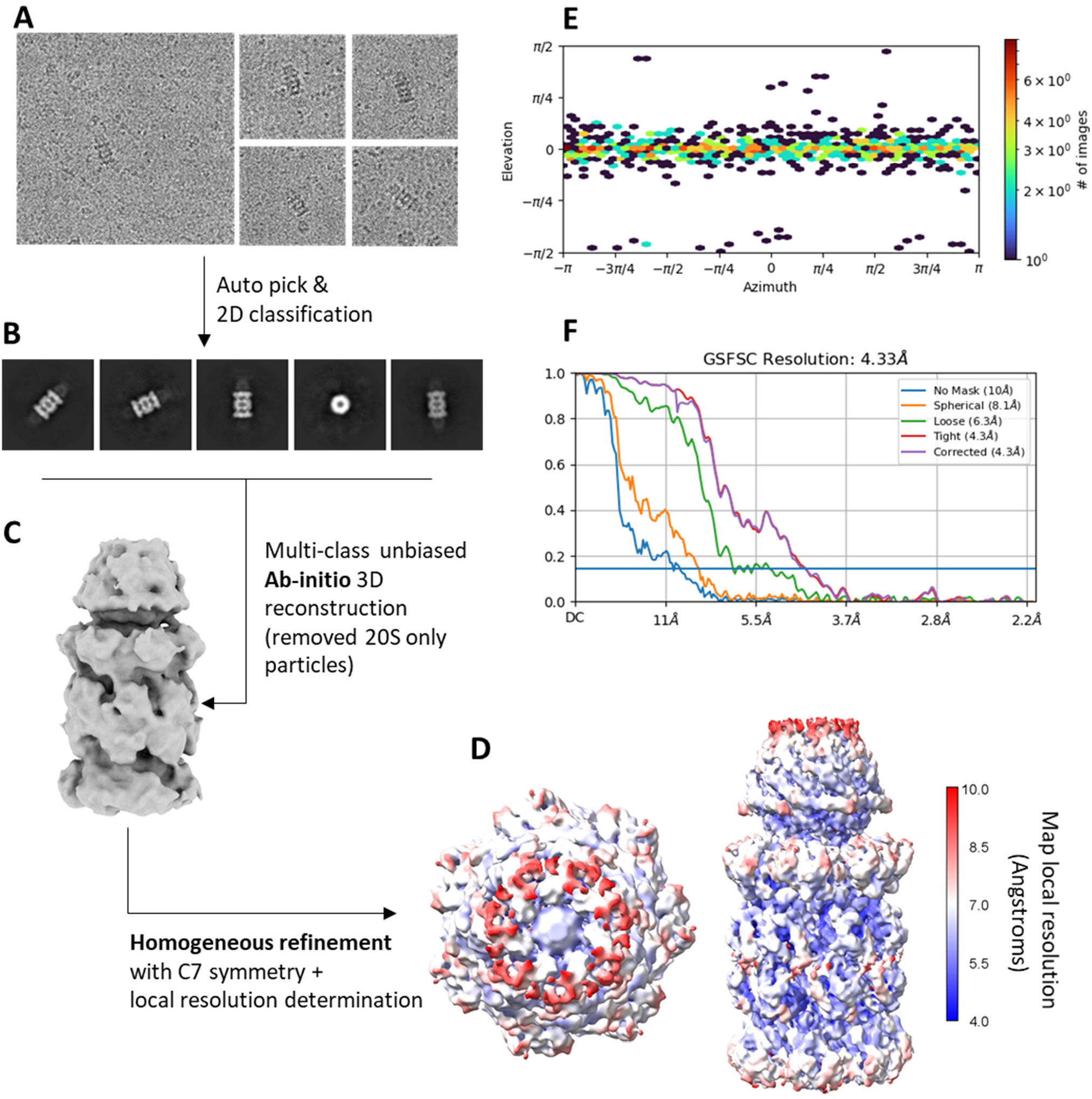
CryoEM workflow and validation. A) Raw motion corrected images of sparsely located PA28γ-20S complexes. B) Several rounds of 2D classification were done to remove junk particles and the final 2D class averages are shown with mostly side views of single and double caped PA28γ-20S complexes, and a single top view. C) Unbiased ab-initio 3D reconstruction with five classes was run to further separate 20S particles from PA28γ-20S complexes (no symmetry was applied at this step). One class contained a clear PA28γ-20S complex (shown) that was derived from 876 particles. D) Particles from the selected ab-initio model were refined with a homogeneous refinement job (with C7 symmetry). The resulting 4.4Å map was colored with outputs from the local resolution job (cryoSPARC) in Chimera to show resolution variability of 3D reconstructed sharpened map. Map resolution colored according to the scale that is shown. E) View of particle angle distribution showing the vast majority of particles that were used for 3D reconstruction were side view. F) Gold standard FSC-0.143 graph showing corrected 4.4Å resolution.

